# Lipid acyl chain remodeling governs KRAS nanoclustering and oncogenic functions through membrane composition control

**DOI:** 10.64898/2026.10.06.757084

**Authors:** Neha Arora, Rafael S. Bravo, Hong Liang, Jeffrey T. Chang, Alex Reed, Junchen Liu, Benjamin F. Cravatt, Haoqiang Ying, Claudia Pedroza, Yong Zhou

**Affiliations:** Department of Diagnostic and Biomedical Sciences, School of Dentistry, University of Texas Health Science Center, Houston, Texas, USA; The Institute for Clinical Research & Learning Health Care, McGovern Medical School, The University of Texas Health Science Center at Houston, Houston, Texas, USA; Department of Integrative Biology and Pharmacology, McGovern Medical School, University of Texas Health Science Center, Houston, Texas, USA; Graduate School of Biological Sciences, MD Anderson Cancer Center and University of Texas Health Science Center, Houston, Texas, USA; Department of Chemistry, The Scripps Research Institute, La Jolla, San Diego, California, USA; Department of Molecular and Cellular Oncology, Division of Basic Science Research, The University of Texas MD Anderson Cancer Center, Houston, Texas, USA

**Keywords:** Lysophosphatidylcholine acyltransferase 3, KRAS, nanoclustering, phosphatidylserine, acyl chains, electron microscopy, pancreatic cancer

## Abstract

KRAS signaling is spatially organized within the dynamic proteolipid nanoclusters on the plasma membrane (PM) that selectively concentrate anionic phospholipids, yet the molecular determinants governing this lipid sorting remain incompletely defined. Here we identify phospholipid acyl chain composition as a critical regulator of KRAS spatial organization and signal output. Using genetic and pharmacological perturbation of lysophosphatidylcholine acyltransferase 3 (LPCAT3), a key enzyme that incorporates polyunsaturated fatty acids into phospholipids, we demonstrate that remodeling of membrane acyl chain unsaturation alters the phosphatidylserine (PS) lipidome without changing total PS abundance. Loss of LPCAT3 depletes polyunsaturated PS species, disrupts KRAS PM association and nanoclustering, and attenuates downstream MAPK signaling and oncogenic activity across multiple KRAS mutant contexts. Acute restoration of polyunsaturated, but not saturated, lipids rescue KRAS nanoclustering, establishing a direct requirement for lipid unsaturation in sustaining signaling-competent assemblies. Mechanistically, these findings define membrane acyl chain composition as a determinant of proteolipid organization that encodes KRAS signaling capacity. Consistent with this model, LPCAT3 inhibition selectively impairs KRAS-dependent proliferation and sensitizes tumor cells to direct KRAS inhibition. Together, this work uncovers lipid acyl chain remodeling as a fundamental regulator of membrane-associated signaling and identifies membrane composition as an exploitable vulnerability in oncogenic KRAS systems.

## Introduction

RAS small GTPases, including isoforms/splice variants KRAS4A, KRAS4B, HRAS and NRAS, are key regulators of essential cell function, such as growth, survival and proliferation. Mutations of *RAS* genes are major drivers of diseases, including cancer, neurodegenerative disorders and cardiovascular diseases. Especially, mutations in *KRAS4B* (hereafter referred to as *KRAS*) are major driving factors of predominantly pancreatic (>90%), colon (45%) and lung (30%) tumors ^1–3^. A hallmark of RAS oncogenesis is spatial compartmentalization of their signaling to distinct proteolipid nanoclusters on the plasma membrane (PM) ^4^. These nanodomains serve as vital signaling platforms to concentrate multiple RAS molecules, receptors, nucleotide exchange factors, effectors and other scaffolds, and to promote efficient signal propagation ^5^. With an averaged diameter of ∼20 nm and lifetime of < 1 second, these nanoclusters are inherently dynamic, permitting RAS-dependent signaling cascades to respond acutely to physical and electrical perturbations typically experienced by cancer cells in various stages of tumor development, including altered cell morphology during epithelial-mesenchymal transition (EMT), metastasis and depolarization of transmembrane potential ^6^. The distinct lipid composition and local membrane environment dictate that the PM is the only efficient signaling compartment for RAS, postioning membrane lipid composition as a regulatory component of RAS signaling.

Significant efforts have focused on understanding how RAS proteins recognize distinct membrane features and respond to membrane perturbations. The most prominent RAS membrane-associating features are their C-terminal lipid anchoring domains ^7^. For KRAS, conformational sampling of its C-terminal farnesylated hexalysine polybasic domain results in its preferred enrichment of anionic phospholipid phosphatidylserine (PS) species with unsaturated *sn-2* acyl chains ^8,9^. We recently reported that the enzymatic globular G-domain of KRAS possesses its own lipid sensing capability, adding another layer of spatial control ^10,11^. Although the G-domains of small GTPases are mostly suspended in the cytosol, they undergo dynamic allosteric reorientation when anchored to the membranes ^12^. Rolling of the G-domains at the membranes results in distinct membrane interfaces ^13^. We discovered that basic residues at these membrane interfaces of KRAS differentially facilitate sensing of lipid acyl chains ^11^. This allosteric-dependent lipid sensing contributes to the allele-specific lipid sensing of KRAS oncogenic mutants ^11^. Although different KRAS oncogenic mutants display allele-specific abilities to enrich saturated lipids, all KRAS mutants tested share enrichment of PS lipids with unsaturated *sn-2* chains ^10,11^. Thus, these findings indicate that RAS proteins possess multilayered, structurally distributed sensitivity to membrane acyl chain architecture within their G-domains and lipid-anchoring domains. Yet the molecular determinants that define this intricate lipid sensing capability remain incompletely understood.

The acyl chain remodeling of phospholipids is primarily catalyzed by a set of enzymes, including elongation of very long-chain fatty acids (ELOVLs), lysophosphatidylcholine acyltransferases (LPCATs) and desaturases. LPCATs use lysophosphatidylcholines (LPCs) and lysophosphatidylethanolamine (LPE) as precursors and various acyl-coAs as donors to directly incorporate fatty acid chains at the *sn-2* position of phospholipids ^14^. In particular, LPCAT3 prefers to incorporate polyunsaturated fatty acid chains at the *sn-2* position of phospholipids, thereby generating polyunsaturated lipids that are selectively enriched in KRAS signaling nanoclusters ^15–17^. It is, thus, possible that LPCAT3 may act as a determinant of KRAS oncogenic signaling.

LPCAT3 has gained attention in cancer research for its role in promoting tumor growth and progression. Specifically, *LPCAT3* expression is higher in tumor tissues versus adjacent tissues in multiple cancers including glioblastoma multiforme (GBM), head and neck squamous cell carcinoma (HNSC), and pancreatic adenocarcinoma (PAAD) ^18^. Blocking LPCAT3 induces cell death in KRAS mutant lung cancer ^18^.

In the current study, we show that genetic and pharmacological perturbations of LPCAT3 remodel PS lipidome, disrupts PM spatial distribution and nanoclustering of KRAS mutants. LPCAT3 downregulation more effectively inhibits proliferation and colony formation of human and mouse pancreatic tumor cell cells dependent on mutant KRAS activities, while having minimal effect on KRAS-independent cells. These data identify LPCAT3-dependent lipid acyl chain remodeling as a mechanistic link between membrane lipid metabolism and RAS nanoscale organization, and establish membrane composition as an active, enzymatically regulated determinant of KRAS signaling.

## Results

### *LPCAT3* knockdown remodels lipidome in MiaPaCa-2 cells

To evaluate the impact of LPCAT3 on remodeling lipid profiles in cells, we generated CRISPR/Cas9 knockdown of *LPCAT3* (*LPCT3*KD) in human pancreatic ductal adenocarcinoma (PDAC) MiaPaCa-2 cells (Fig.1A). RNA sequencing validated the significant decrease in *LPCAT3* expression in MiaPaCa-2 cells with *LPCAT3*KD when compared to the vector control (V2) (Fig.1B). When compared with the vector control, significantly downregulated genes in the *LPCAT3*KD MiaPaCa-2 cells included those associated with PS lipid metabolism and small GTPase-mediated signaling, such as *GPAT3 (*glycerol-3-phosphate acyltransferase 3)*, ALDH1A* (Aldehyde Dehydrogenase 1)*, SERPINF1* (serpin family F member 1), *RCAN1* (Regulator of Calcineurin 1), *ARHGAP29* (Rho GTPase-activating protein 29) (Fig.1C-E). Correspondingly, pathway enrichment analysis showed significant decrease in PS metabolic process, small GTPase-mediated signal transduction, regulation of small GTPase-mediated signal transduction and epithelial cell proliferation in the *LPCAT3*KD MiaPaCa-2 cells (Fig.1D).

**Figure 1.**
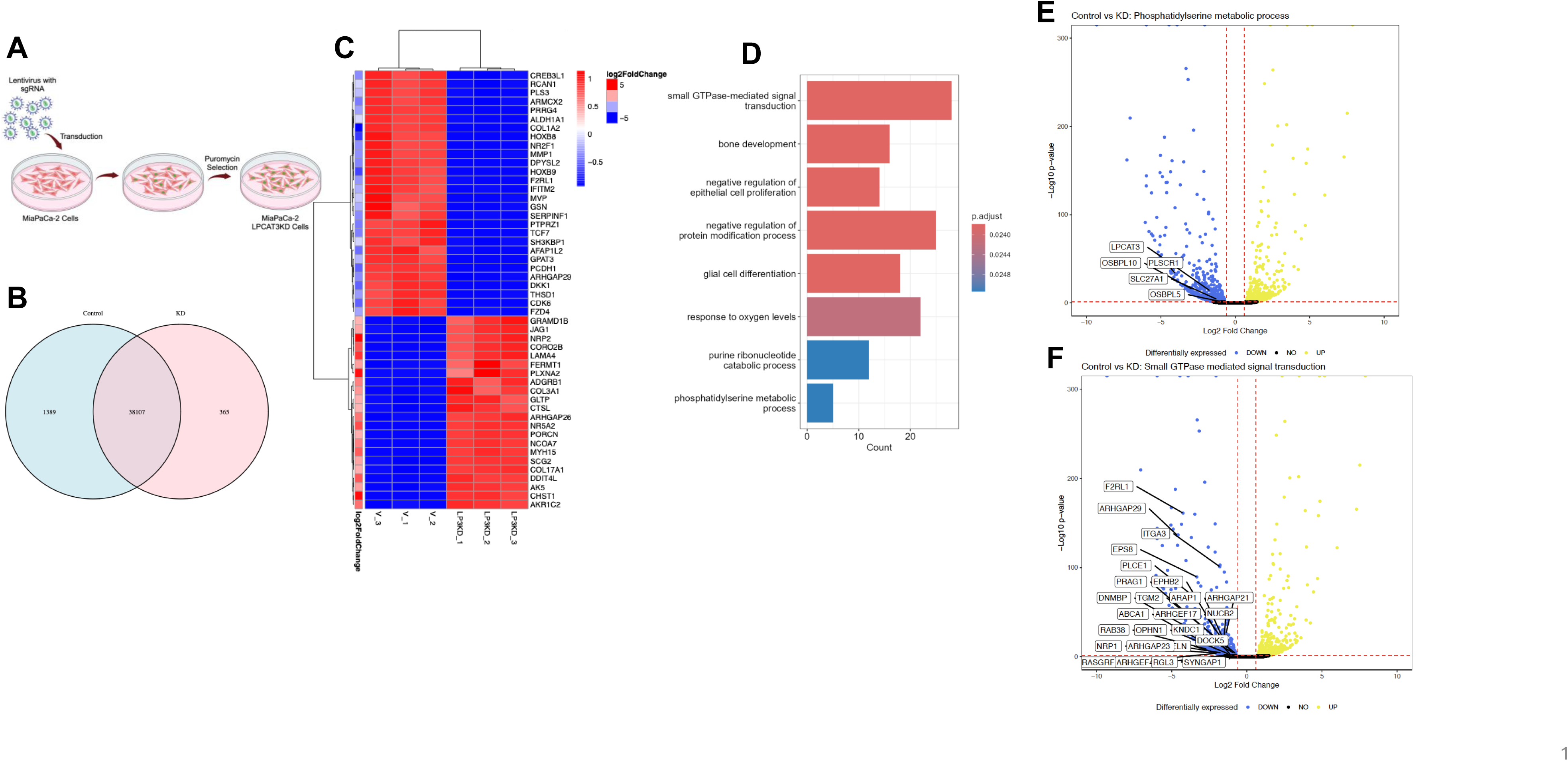
LPCAT3 deletion remodels lipid metabolism. Human PDAC MiaPaCa-2 cells expressing vector control (V2) or sgRNA of *LPCAT3* (*LPCAT3*KD) were grown to a monolayer before harvesting for RNA sequencing (A), which validated reduction of *LPCAT3* RNA expression (B). (C) A heat map shows genes markedly altered MiaPaCa-2 cells harboring V2 or *LPCAT3*KD. (D) Pathway enrichment analysis identifies biological pathways disrupted in LPCAT3KD cells when compared with the vector control. (E and F) Volcano plots identify the reduced gene expression in networks of small GTPase signaling and lipid metabolism. Data shown includes three individual trials for each condition. Three individual trials were conducted and analyzed for each condition.

We next compared lipid profiles of MiaPaca-2 cells with vector control and *LPCAT3*KD in shotgun lipidomics. MiaPaCa-2 *LPCAT3*KD cells displayed distinct lipid profiles than those expressing vector control (Fig.2A), with many polyunsaturated phosphatidylcholine (PC), phosphatidylethanolamine (PE) and phosphatidylserine (PS), phosphoinositols (PI), cholesterol esters (ChE) and triacylglycerides (TG) species altered (Fig.2B-H). Interestingly, while total PE and PS levels remained similar, total PC level was lower in *LPCAT3*KD cells than the vector control (Fig.2I-K). C18:2 CoA and C20:4 CoA are the most preferred ligands of LPCAT3 ^19^. In Supplemental Fig.S1, LPCAT3 deletion had various effects on homeostasis of major lipids species with 18:2 or 20:4 *sn-2* chains, with the most significant reduction of PE species with 20:4 *sn-2* chains. Since PS lipids are key structural components of KRAS signaling scaffolds ^5,6^, we next compared the PS profiles. Total PS levels were not altered in *LPCAT3*KD cells when compared with vector control (Fig.2K). Major PS species, including 16:0/16:1 PS, 18:0/20:2 PS, 18:1/22:1 PS and 22:0/18:2 PS, were reduced in MiaPaCa-2 cells with *LPCAT3*KD when compared with the vector control (Fig.3A-I). Some minor PS species, such as 18:1/20:3 and 20:4/18:2 PS, were elevated in cells with *LPCAT3*KD. Taken together, *LPCAT3* deletion remodels lipidome of human PDAC cells, especially depleting major polyunsaturated PS species.

**Figure 2.**
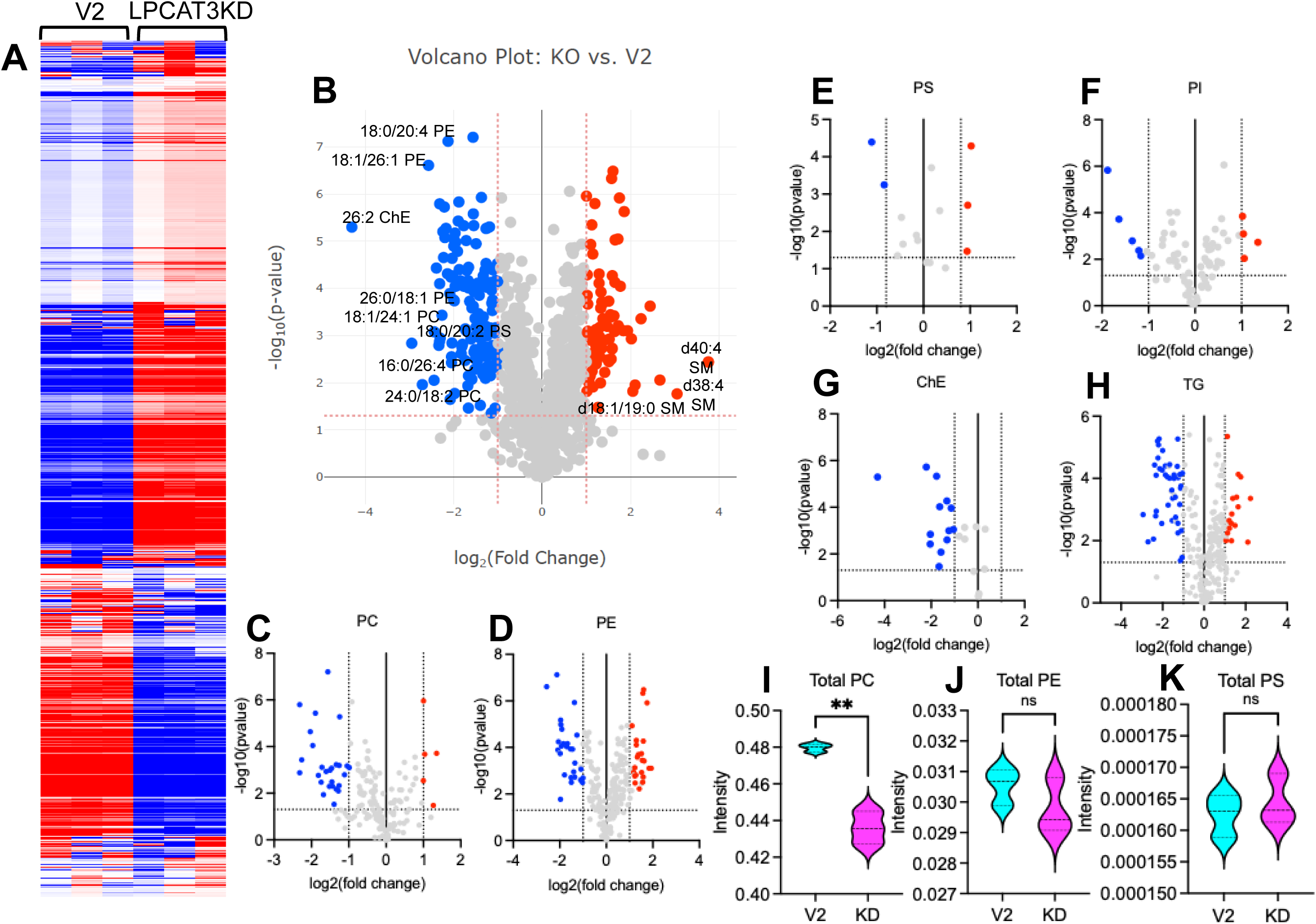
*LPCAT3* deletion modulates lipidome of human PDAC MiaPaCa-2 cells. (**A**) A heatmap summarizes altered lipidomes of MiaPaCa-2 cells expressing vector control or sgRNA of *LPCAT3*. Volcano plots illustrate the most decreased and elevated species of global lipidomes (B), PC (C), PS (D), cholesterol esters (ChE, E), PE (F), PI (G), triacylglycerides (TG, H) in MiaPaCa-2 cells expressing vector control and *LPCAT3*KD. Violin plots of total PC (I), total PE (J), total PS (K) are shown. Data includes three individual trials for each condition.

**Figure 3.**
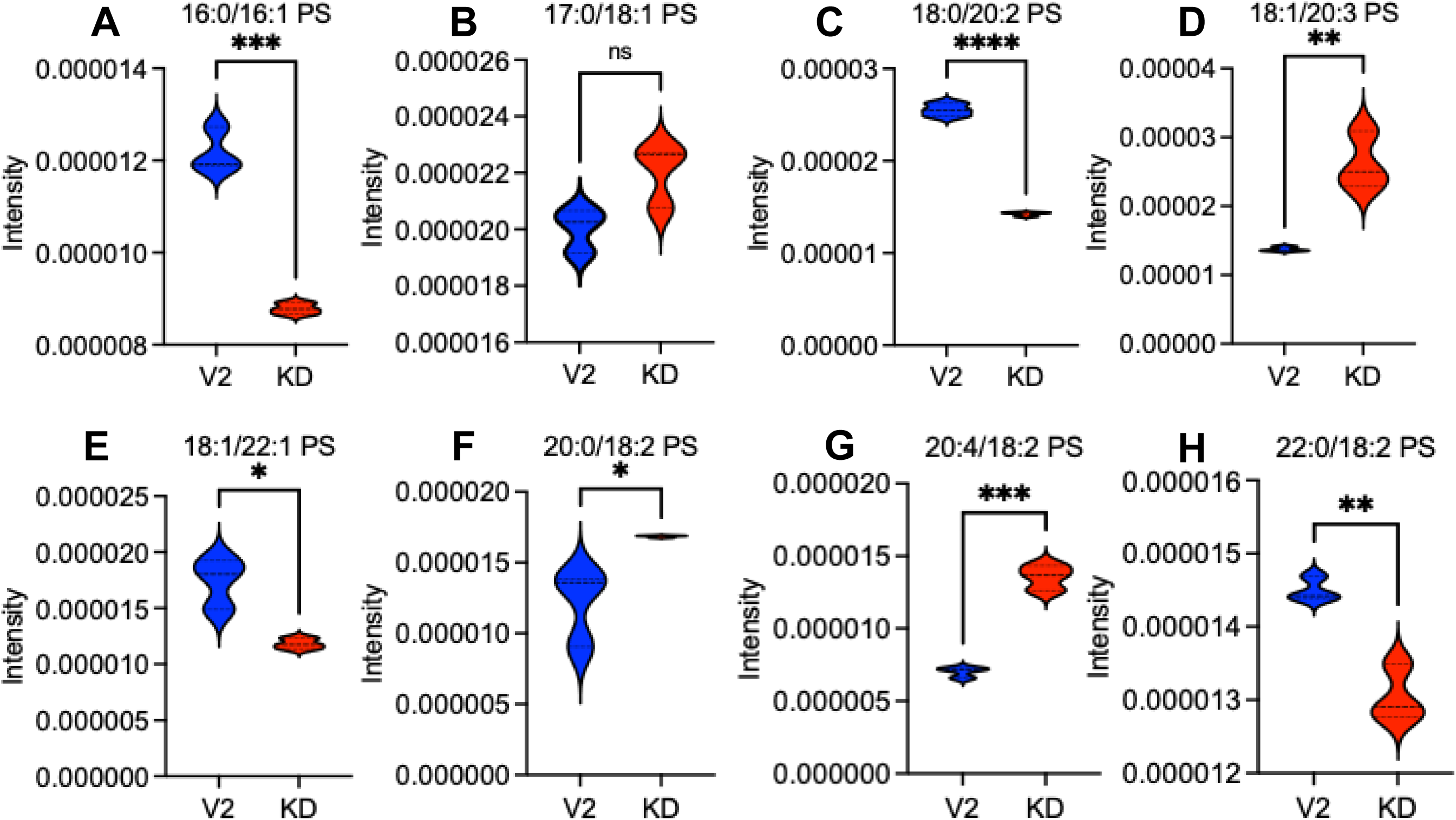
LPCAT3 deletion remodels homeostasis of PS lipids. Violin plots summarize changes in specific PS species, such as 16:0/16:1 PS (A), 17:0/18:1 PS (B), 18:0/20:2 PS (C), 18:1/20:3 PS (D), 18:1/22:1 PS (E), 20:0/18:2 PS (F), 20:4/18:2 PS (G) and 22:0/18:2 PS (H). Data includes three individual trials for each condition. Two-tail unpaired t-test was used to evaluate statistical significance between V2 and LPCAT3KD groups, * indicating p < 0.05, ** p < 0.01, *** p < 0.001, **** < 0.0001.

### LPCAT3 downregulation disrupts spatial distribution of mutant KRAS on the plasma membrane

Unsaturated PS lipids are critical structural components of KRAS signaling nanoclusters on the plasma membrane (PM) and play key roles in facilitating proper and efficient KRAS intracellular transport and formation of signaling scaffolds ^5,20,21^. To examine how LPCAT3 downregulation impacted the spatial distribution of KRAS mutants, we performed electron microscopy (EM)-univariate nanoclustering analysis ^10,11,22,23^. Briefly, MiaPaCa-2 cells with vector control or *LPCAT3*KD were grown on gold EM grids and were transfected with *GFP-KRAS^G12V^*. Intact basal PM sheets attached to the EM grids were immunolabeled with anti-GFP antibody conjugated to 4.5-nm gold nanoparticles. Following transmission EM (TEM) imaging at a magnification of 100,000X and digitization, spatial distribution of the gold-labeled GFP-KRAS^G12V^ within a select 1 μm^2^ PM area was calculated using the Ripley’s univariate K-function (Supplemental Fig.2A and B). Extent of nanoclustering, *L*(*r*) – *r*, was plotted as a function of length scale *r* (Supplemental Fig.2C). The peak *L*(*r*) – *r* value, termed as Lmax (Supplemental Fig.2C), was used as a statistical summary for nanoclustering. *L*(*r*) – *r* value of 1 is the 99% confidence interval (99% CI), values above which indicate statistical meaningful nanoclustering (Supplemental Fig.2C). Total gold number per 1 μm^2^ PM area was counted to indicate PM localization. We found that Lmax of GFP-KRAS^G12V^ (Fig.4A) and gold labeling (Fig.4B) were significantly reduced in *LPCAT3*KD MiaPaCa-2 cells when compared with the vector control, suggesting that *LPCAT3* deletion mislocalizes KRAS^G12V^ from the PM and disrupts the nanoclustering of KRAS^G12V^ left on the PM. In a rescue experiment, exogenous PS species were acutely added back (10 μM, 1 hour) to the *LPCAT3*KD MiaPaCa-2 cells. Acute addback of natural extracts of mouse brain PS enriched with polyunsaturated PS species, but not saturated di18:0 PS (DSPS), effectively restored the nanoclustering and PM localization of GFP-KRAS^G12V^ (Fig.4A and B). We further interrogated our spatial analysis data to obtain effects of *LPCAT3* deletion on population distribution of KRAS molecules on the PM. *LPCAT3* deletion significantly increased monomer population and reduced trimer and higher-order oligomer populations of GFP-KRAS^G12V^ (Supplemental Fig.S2E-H). We next tested pharmacological inhibition of LPCAT3, using a recently developed LPCAT3 inhibitor (R)-HTS3 ^19^. MiaPaCa-2 cells ectopically expressing *GFP-KRAS^G12V^* were treated with DMSO or 10 μM HTS3 for 24 hours before EM-univariate analysis. HTS3 treatment effectively mislocalized GFP-KRAS^G12V^ from the PM (Fig.4C) and disrupted the nanoclustering of GFP-KRAS^G12V^ left on the PM (Fig.4D). Taken together, genetic and pharmacological downregulation of LPCAT3 disrupts the PM association of mutant KRAS.

**Figure 4.**
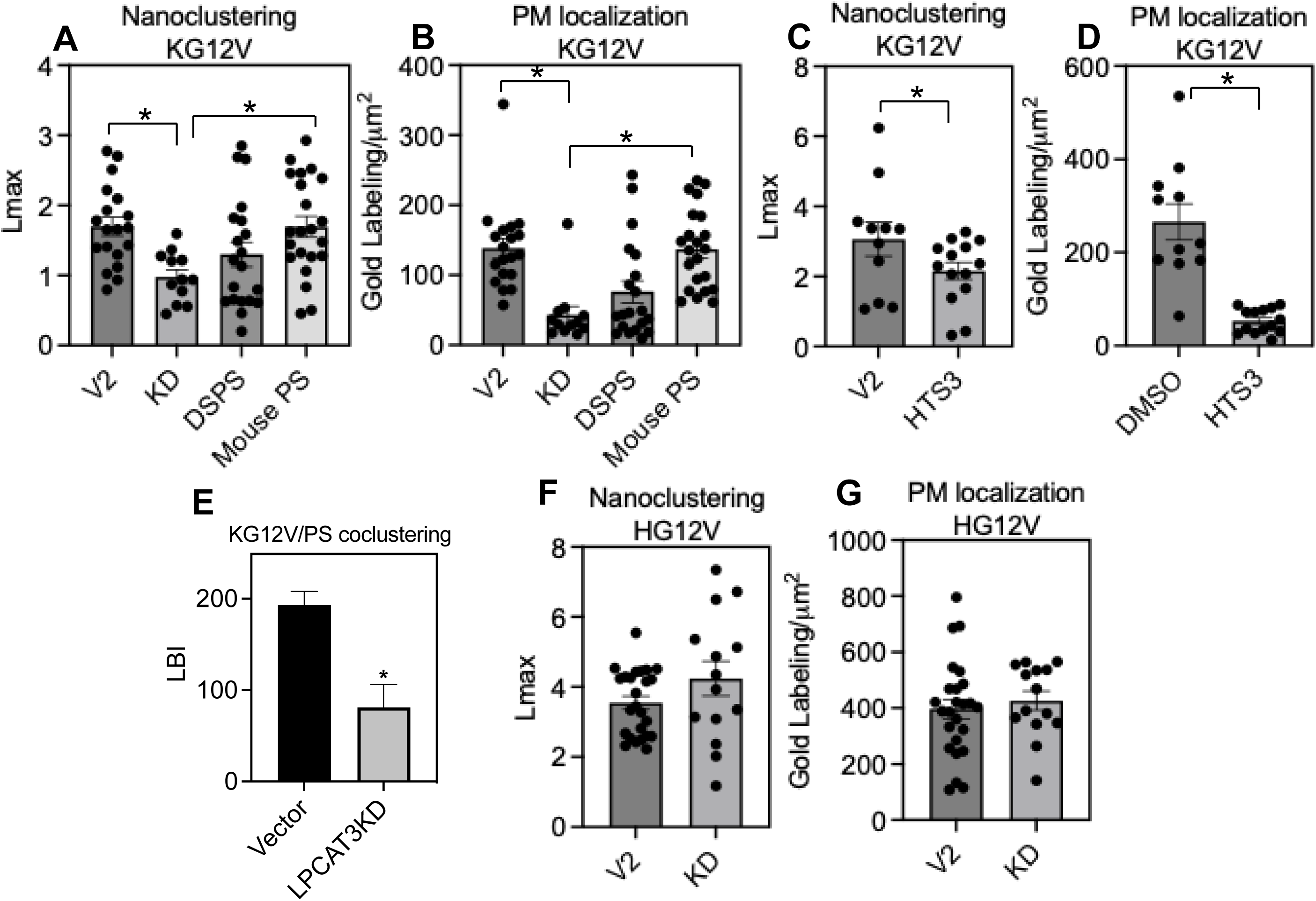
*LPCAT3* deletion selectively disrupts the plasma membrane spatial distribution of mutant KRAS. Human PDAC MiaPaCa-2 cells stably expressing vector control or *Lpcat3*KD were transiently transfected with *GFP-KRAS^G12V^*. GFP-KRAS^G12V^ anchored to the intact PM sheets of these cells were immunolabeled with 4.5 nm nanoparticles conjugated to anti-GFP antibody. A representative EM image is illustrated in supplemental Fig. S2. Distribution of the gold-labeled GFP-KRAS^G12V^ within a selected 1μm^2^ PM area was calculated using the Ripley’s K-function analysis. A nanoclustering curve was plotted as the extent of nanoclustering, *L*(*r*) – *r*, vs. length scale, *r* in nanometers. The peak value of the curve, termed as *L_max_*, was used as a summary statistic to indicate nanoclustering (A). Total gold labeling with the same 1μm^2^ PM area indicates extent of PM localization (B). To rescue effects of *Lpcat3* KD on spatial distribution of KRAS^G12V^, we performed lipid acute addback experiments, where MiaPaCa-2 cells were incubated with exogenous PS species (DSPS, or mouse brain PS at 10 μM) for 1 hour before EM spatial analysis. To test pharmacological inhibition of LPCAT3, Vector control and *LPCAT3*KD MiaPaCao-2 cells transiently expressing GFP-KRAS^G12V^ were treated with DMSO or LPCAT3 inhibitor (R)-HTS3 (10 μM, 48 hours). EM spatial analysis calculated extent of nanoclustering (C) and gold labeling (D) of GFP-KRAS^G12V^ in these cells. (E) To quantify effects of LPCAT3 on PS enrichment of KRAS, vector control and *LPCAT3*KD MiaPaCao-2 cells transiently expressing GFP-LactC2 (a PS-specific domain) and RFP-KRAS^G12V^ were used. Intact PM sheets were immunolabeled with 6 nm gold particles conjugated to anti-GFP antibody and 2 nm gold coupled with anti-RFP antibody. The Ripley’s bivariate co-clustering K-function calculated extent of co-localization between PS and KRAS. Similar EM spatial analysis quantified extent of nanoclustering (F) and gold labeling (G) of GFP-HRAS^G12V^ in MiaPaCa-2 cells with vector control or *LPCAT3*KD. At least 15 EM images from individual cells were imaged and analyzed for each condition. Data are shown as mean ± SEM. For the gold labeling data, the statistical significance was quantified using the one-way ANOVA. For the univariate nanoclustering and bivariate co-clustering data, the statistical significance was evaluated via the non-parametric bootstrap tests. * Indicates p < 0.05.

To examine whether the reduced KRAS PM association was caused by the disrupted PS enrichment in KRAS nanoclusters, we used bivariate K-function analysis to quantify the co-clustering of PS lipids and KRAS in MiaPaCa-2 cells. Vector control and *LPCAT3*KD MiaPaCa-2 cells were co-transfected with *RFP-KRAS^G12V^*and *GFP-LactC2*, a specific PS-binding domain. Intact basaolateral PM sheets of MiaPaCa-2 cells on the EM grids were immunolabeled with 6 nm gold nanoparticles conjugated to anti-GFP antibody and 2 nm gold linked to anti-RFP antibody, respectively. Following TEM imaging, co-clustering between the two gold populations was calculated via bivariate K-Function analysis. Extent of co-clustering, *L_biv_*(*r*) – *r*, was plotted as a function of length scale *r* (Supplemental Fig.2D). Area-under-the-curve of *L_biv_*(*r*) – *r* vs. *r* between r=10 and 110 was termed as L-function bivariate integrated (LBI) and used as a statistical summary for the co-clustering. The LBI value of 100 is the 95% confidence interval (95% CI), values above which indicate statistical meaningful co-clustering. Fig.4E shows *LPCAT3* deletion significantly reduced the co-clustering of KRAS^G12V^ and PS, thus depleting PS from KRAS nanoclusters. Taken together, *LPCAT3* deletion disrupts PS enrichment of KRAS^G12V^ on the PM. We and others previously showed that the PM association of an oncogenic mutant of another RAS isoform, HRAS^G12V^, is independent of PS ^8,22,24,25^. To examine specificity of LPCAT3 downregulation, we measured effects of *LPCAT3* deletion on the nanoclustering of GFP-HRAS^G12V^ in MiaPaCa-2 cells with vector control or *LPCAT3*KD. *LPCAT3* deletion had no effect on the nanoclustering (Fig.4F) and PM localization (Fig.4G) of GFP-HRAS^G12V^. Taken together, *LPCAT3* downregulation had no effect on spatial organization of HRAS.

Our data suggests that the membrane core properties defined by lipid acyl chain structures are critical determinants of KRAS nanoclustering. The LPCAT3-synthesized polyunsaturated lipids are key susceptible substrates of ferroptosis, a new form of non-apoptotic and iron-dependent programmed cell death triggered by the accumulation of peroxidized lipids ^26–29^. KRAS and ferroptosis are closely intertwined, with RAS (including KRAS)-transformed tumor cells being especially sensitive to ferroptosis ^30–32^. It is, thus, possible that the observed effects of LPCAT3 downregulation on KRAS nanoclustering may be in part a result of perturbed ferroptosis. To test this, we treated BHK cells with a small molecule inducer of ferroptosis, RAS-selective lethal 3 (RSL3). RSL3 deactivates glutathione peroxidase 4 (GPX4), an enzyme that eliminates lipid peroxides, thus causing accumulation of peroxidized lipids and triggering ferroptosis. RSL3 treatment had no effect on the PM localization (Supplemental Fig.S3A) and nanoclustering (Supplemental Fig.S3B) of GFP-KRAS^G12V^, as well as the co-clustering of PS and KRAS^G12V^ (Supplemental Fig.S3C). Interestingly, RSL3 treatment disrupted the nanoclustering (Supplemental Fig.S3D), but promoted the PM localization (Supplemental Fig.S3E), of PS lipids in BHK cells. EM-bivariate co-clustering analysis illustrates that RSL3 treatment disrupted co-localization between PS and cholesterol in BHK cells (Supplemental Fig.3F). Thus, the induced ferroptosis alters spatial distribution of PS lipids, without directing impacting that of KRAS. Taken together, genetic and pharmacological downregulation of LPCAT3 selectively disrupts spatial organization of a KRAS oncogenic mutant, without affecting that of an HRAS mutant. This is likely a direct effect of the altered polyunsaturated lipids and unrelated to ferroptosis.

### Oncogenic activities of mutant KRAS depend on LPCAT3

To compare effects of *LPCAT3* deletion on oncogenic activities of KRAS mutants, MiaPaCa-2 cells with vector control or *LPCAT3*KD were grown for 72 hours for proliferation assay and 10 days for colony formation. *LPCAT3* deletion significantly reduced proliferation (Fig.5A) and colony formation (Fig.5B) of MiaPaCa-2 cells. Anchorage-independent growth of cancer cells is a key aspect of the tumor phenotype and stringent assessment of tumorigenic properties. MiaPaCa-2 *LPCAT3*KD cells showed significantly reduced number of colonies in soft agar assay when compared with the vector control, as determined at week 3 after incubation (Fig.5C). Taken Together, *LPCAT3* deletion compromises the tumorigenic properties of human PDAC MiaPaCa-2 cells.

**Figure 5.**
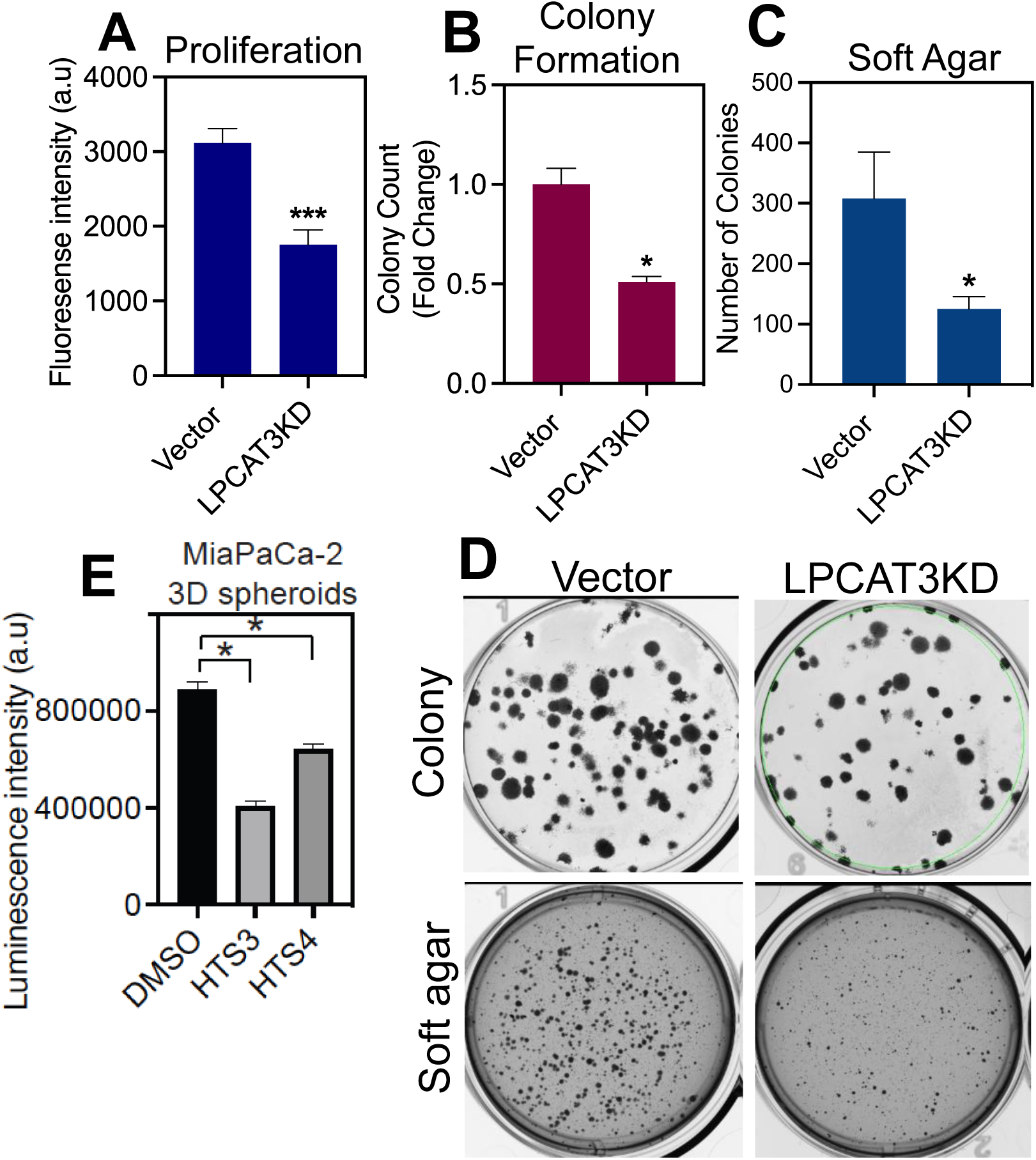
*LPCAT3* deletion reduces oncogenic activities of human PDAC cells. (A) MiaPaCa-2 vector control or *LPCAT3*KD cells were seeded in 96-well black plate and grown for 72 hours. Cell numbers were determined by CyQuant Proliferation Assay. (B) MiaPaCa-2 vector control and *LPCAT3*KD cells were seeded in 6-well plate. After 10 days of growth, cell colonies were stained with 0.01% crystal violet and imaged. Colony numbers were quantified using ImageJ. (C) MiaPaCa-2 vector control and *LPCAT3*KD cells were seeded in soft agar, with a base layer of 1% agar–media mixture and a top layer of a 0.6% agar–cell suspension in six-well plates. After 3 weeks, cell colonies were stained with 0.01% crystal violet and imaged. Colony numbers were quantified using ImageJ. For all assays, 3 individual trials were conducted and data shown as mean ± SEM. Statistical significance was evaluated using *t* tests (*p<0.05).

### KRAS-dependent tumor cells are more sensitive to LPCAT3 downregulation

To investigate specificity of LPCAT3 deletion on KRAS oncogenesis, we tested a murine pancreatic cancer iKRAS AK192 cell line with doxycycline (DOX)-induced expression of KRAS^G12D^, thereafter termed as iKRAS cells ^33,34^. Fig.6A shows DOX treatment effectively upregulated phosphorylation of ERK (pERK/total ERK, a readout of KRAS-MAPK signal output), confirming the functionality of the inducible system. Three siRNA constructs were directed against *Lpcat3* in murine iKRAS cells. Fig.6B shows a significant decrease in LPCAT3 protein expression upon treatment with siRNA1 or siRNA2 as compared to the scramble control siRNA in Western blotting. Expression of siRNA1 or siRNA2 reduced pERK/total ERK levels in iKRAS cells induced to express KRAS^G12D^ (DOX+), while having no effect on MAPK signaling in cells without KRAS^G12D^ expression (DOX-) (Fig.6C and D). Consistently, treatment with siRNA1 or siRNA2 against *Lpcat3* significantly reduced colony formation of iKRAS cells induced to express KRAS^G12D^ (DOX+), while having no effect in the uninduced condition (DOX-, Fig.6E-G). Taken together, *Lpcat3* deletion inhibits oncogenic activities of mutant KRAS-dependent tumor cells, without affecting KRAS-independent cells.

**Figure 6.**
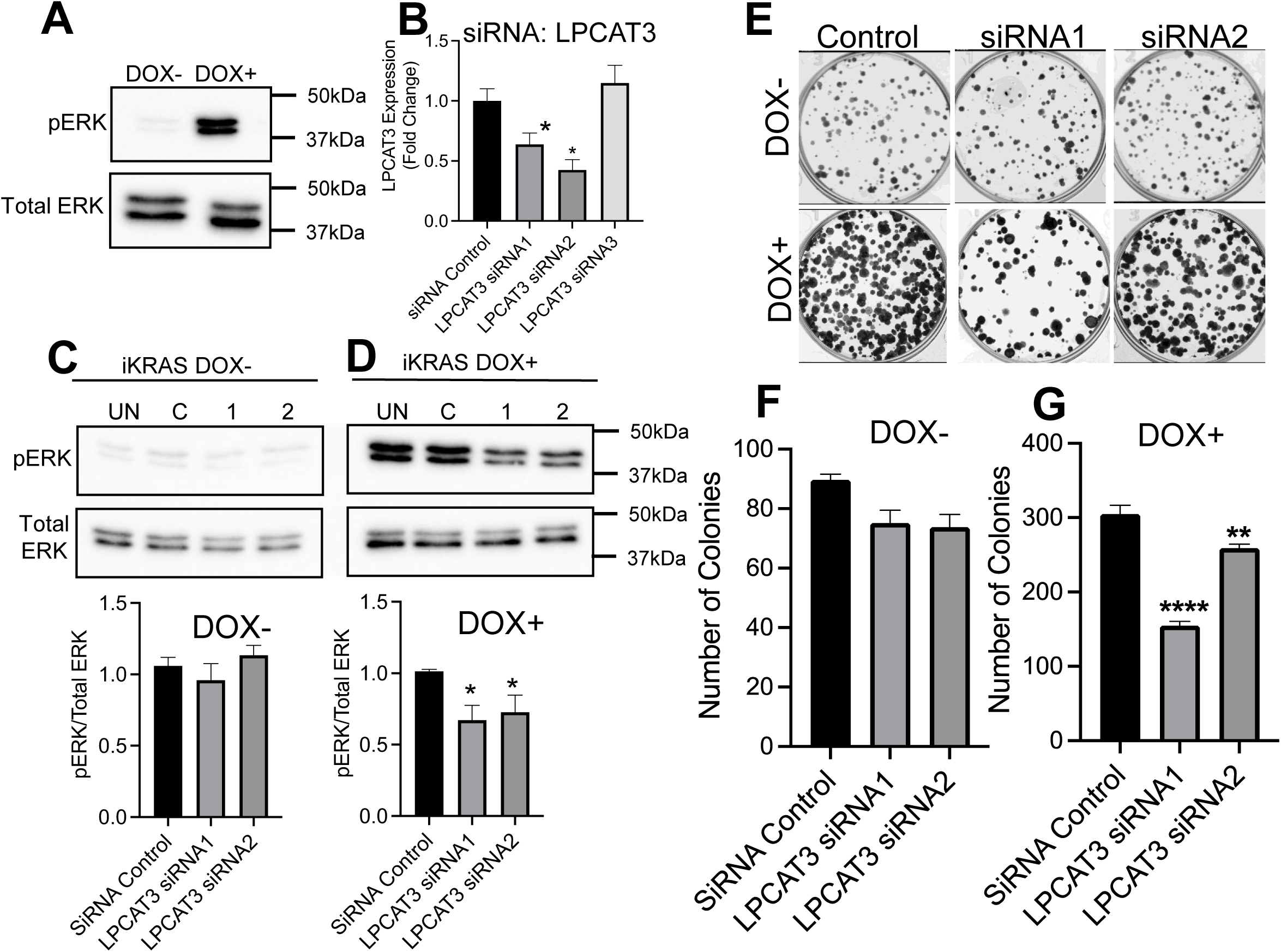
KRAS-dependent tumor cells are more sensitive to *Lpcat3* knockdown. (A) Whole cell lysate of mouse PDAC iKRAS cells were collected and subjected to western blot analysis. Western blot analysis of LPCAT3 expression levels was performed to validate siRNA mediated *Lpcat3*KD in iKRAS cells. iKRAS (DOX+) and iKRAS (DOX-) cells treated with 10 nM siRNA targeting *Lpcat3*. Blot quantification was performed using ImageJ and statistical analysis was performed using one way ANOVA (*p<0.05). (B) Western blot analysis to validate induction of MAPK signaling measured by pERK levels in DOX+ iKRAS cells. (C) Western blot analysis of pERK levels in iKRAS cells with or without DOX after treatment with *Lpcat3* siRNA. Blot quantification was performed using ImageJ and statistical analysis was performed using one way ANOVA (*p<0.05). (D) iKRAS cells with or without DOX were seeded in 6-well plate. Following attachment, cells were treated with *Lpcat3* siRNA. After 10 days, the colonies were stained with 0.01% crystal violet and imaged. Colony numbers were quantified by ImageJ. Three individual trials were conducted for each condition. Data are shown as mean ± SEM, and significant differences were evaluated using *t* tests (* p<0.05).

We next explored specific effects of LPCAT3 inhibitors on KRAS oncogenic activities. Human PDAC *KRAS^G12C^*-expressing MiaPaCa-2 cells and wild-type *KRAS*-expressing BxPC-3 cells were treated with DMSO or LPCAT3 inhibitors (R)-HTS3 or HTS4 before measuring MAPK signal output. In Fig.7A-C, treatment of (R)-HTS3 or HTS4 significantly reduced levels of pERK/total ERK in MiaPaCa-2 cells, while having no effect on those of BxPC-3 cells. Using the inducible expression system, we next show that treatment of (R)-HTS3 or HTS4 compromised colony formation of iKRAS cells induced to express *KRAS^G12D^* (DOX+), while having no effect on iKRAS cells without mutant *KRAS* expression (DOX-, Fig.7D-F). In MiaPaCa-2 cells, (R)-HTS3, but not HTS4, statistically significantly reduced colony formation of MiaPaCa-2 cells (Fig.7G). Taken together, genetic and pharmacological downregulation of LPCAT3 selectively compromises signaling and oncogenic activities of mutant KRAS-dependent tumor cells.

**Figure 7.**
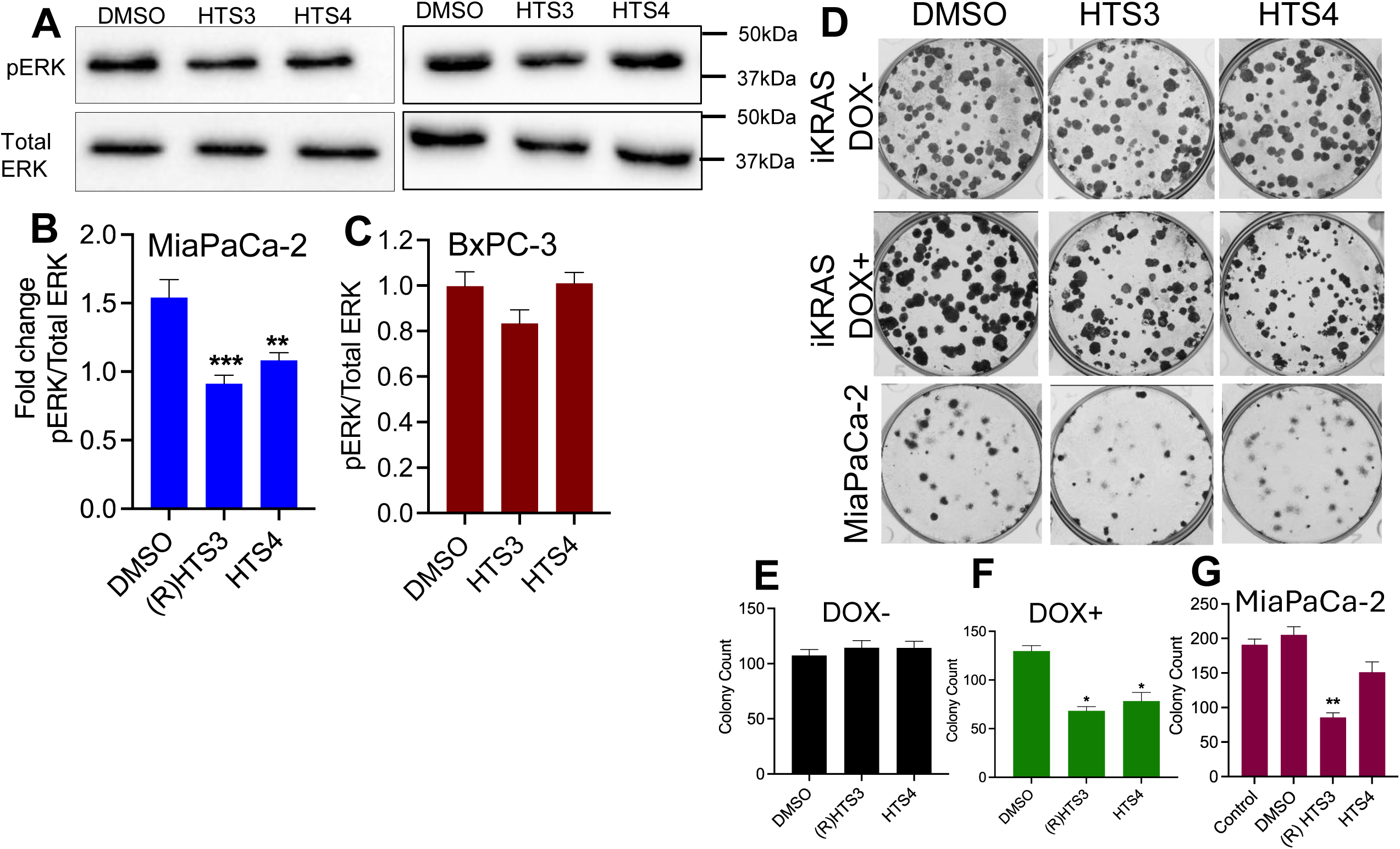
LPCAT3 inhibition compromises KRAS-dependent oncogenic functions. (A) Whole cell lysate from MiaPaCa-2 and BxPC-3 cells treated with vehicle DMSO, 10 μM of (R)HTS3 or HTS4 for 72 h were subjected to western blot analysis to evaluate pERK and total ERK expression levels. Blot quantification was performed using ImageJ and statistical analysis was performed using one way ANOVA (*p<0.05). (B) iKRAS cells with or without DOX and MiaPaCa-2 cells were seeded in 6-well plate. Following attachment, cells were treated with DMSO, 10 μM of (R)HTS3 or HTS4, colonies were stained with 0.01% crystal violet and imaged. Colony numbers were quantified by ImageJ. Three individual trials were conducted for each condition. Data are shown as mean ± SEM, and significant differences were evaluated using *t* tests (* p<0.05).

### LPCAT3 inhibition suppresses oncogenic activitity of multiple KRAS mutants

Signaling nanoclusters of different KRAS mutants share unsaturated PS lipids ^11^, thus potentially susceptible to LPCAT3 deletion. To evaluate effects of LPCAT3 inhibition on oncogenic activity of different KRAS mutants, we compared human PDAC lines including MiaPaca-2 (*KRAS^G12C^*), MoH (*KRAS^G12R^*), AsPC-1 (*KRAS^G12D^*), Capan-1 (*KRAS^G12V^*) and Patu8902S (*KRAS^G12V^*), as well as PDAC BxPC-3 cells and immortalized pancreatic epithelial HPNE cells both of which express wild-type *KRAS*. These human pancreatic cell lines were treated with DMSO, (R)-HTS3 or HTS4 for 72 hours at different doses, followed by measuring cell viability. (R) HTS3 significantly reduced viability of human PDAC cells bearing KRAS mutation and had no significant effect on BxPC-3 and HPNE cells (Fig.8 A-G). HTS4 had similar specific effects on KRAS mutatnt-exppressing PDAC cells, except for MOH and Capan-1 cells (Fig.8H-N). Taken together, *KRAS*-dependent PDAC cells are more sensitive to LPCAT3 inhibition than *KRAS*-independent cells.

**Figure 8.**
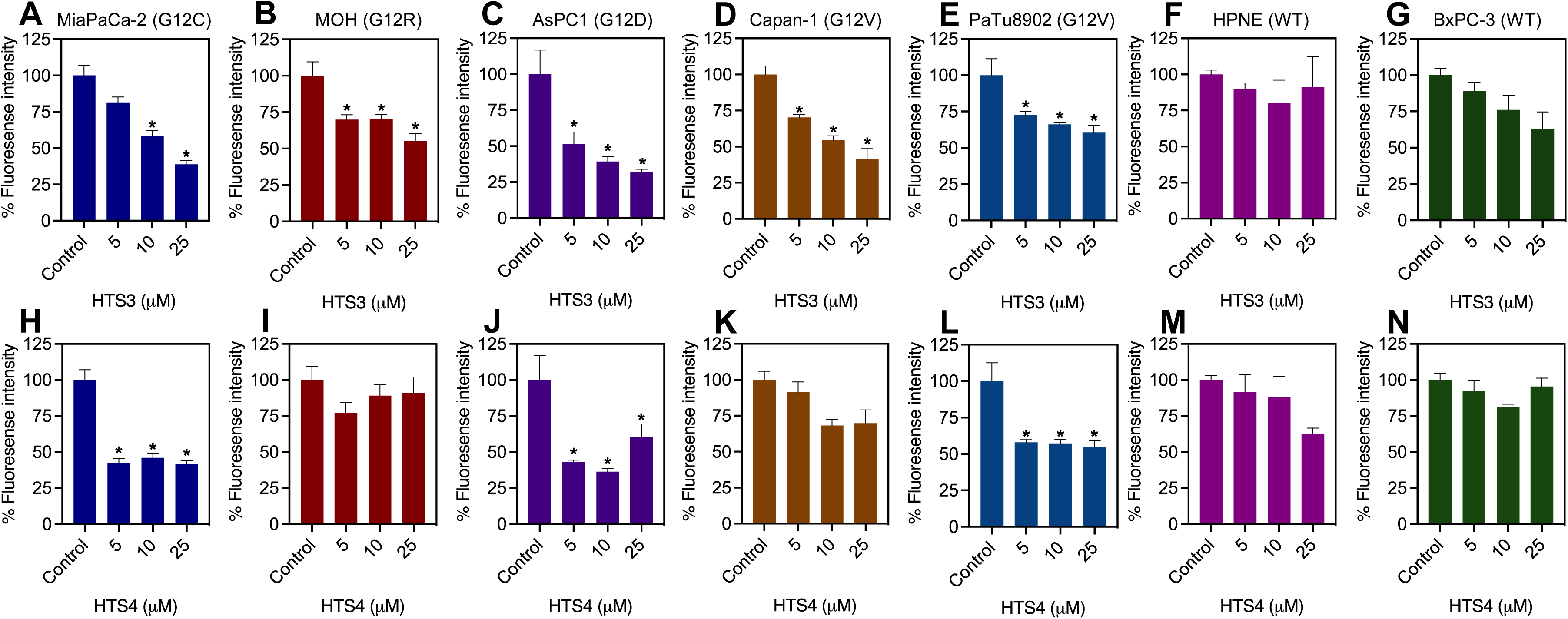
LPCAT3 inhibition suppresses proliferation of human PDAC cell lines. Human PDAC lines harboring different. KRAS mutants were treated with DMSO, 10 μM of (R)HTS3 or HTS4 for 72 hours before CyQuant assay to measure proliferation. Cell lines include MiaPaCaq-2 (G12C, A and H), MOH (G12R, B and I), AsPC1 (G12D, C and J), Capan-1 (G12V, D and K), PaTu8902 (G12V, E and L), HPNE (wild-type, F and M) and BxPC-3 (wild-type, G and N). Three indivudal trials were conducted for each condition. Data are shown as mean ± SEM, and significant differences were evaluated using *t* tests (* p<0.05).

**Figure 9.**
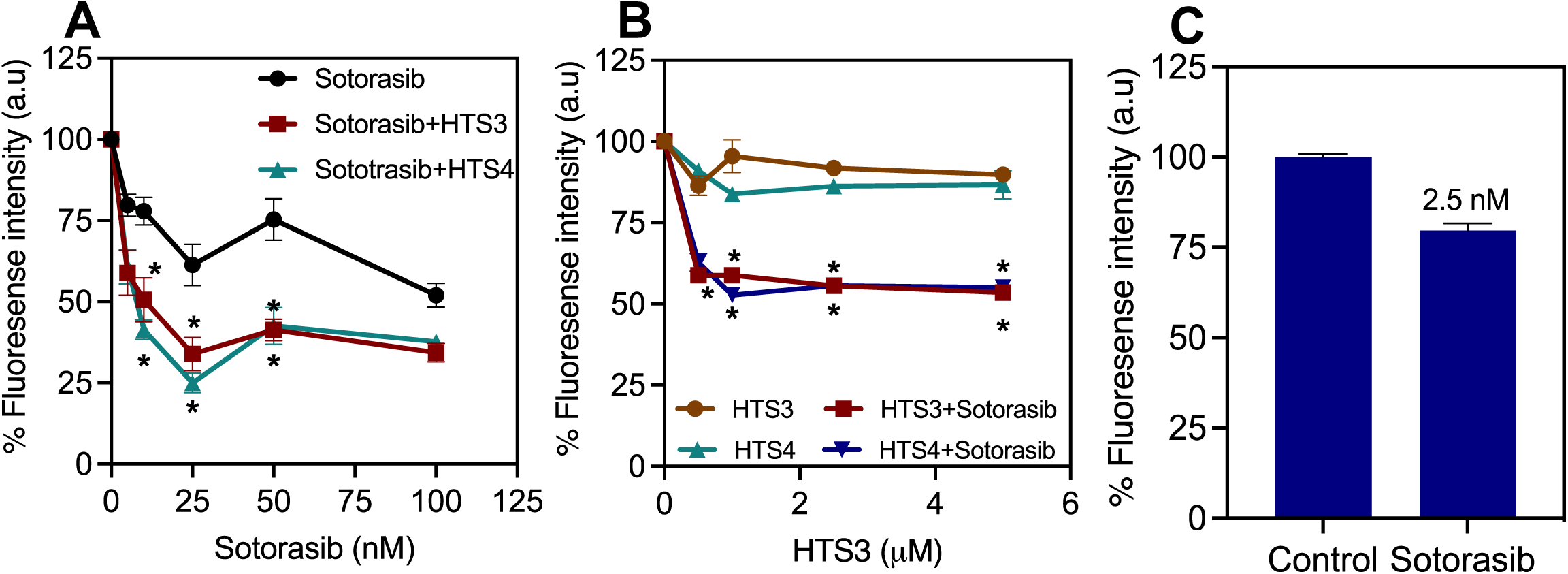
LPCAT3 inhibition sensitizes human PDAC cells to KRAS direct inhibitor. (A) MiaPaCa-2 cells were seeded in 96 well plates and treated with a fixed sub-optimal 5 μM concentration of (R)HTS3 or HTS4 without/with varying concentration of Sotorasib. Following 72 h of incubation, cell numbers were determined by CyQuant Proliferation Assay. (B) MiaPaCa-2 cells were seeded in 96 well plates and treated with a fixed sub-optimal 2.5 nM Sotorasib without/with varying concentrations of (R)HTS3 or HTS4. Following 72 h of treatment, cell numbers were determined by CyQuant Proliferation Assay. Data are shown as mean ± SEM. Statistical significance was evaluated using Two-way ANOVA (*p < 0.05).

### LPCAT3 inhibition sensitizes human PDAC cells to direct KRAS inhibitor

Targeting LPCAT3 impacts the lipid-dependent spatiotemporal organization of mutant KRAS, while KRAS direct inhibitors bind to the enzymatic G-domains of KRAS mutants. It is, thus, possible that combined inhibition of LPCAT3 and KRAS may be a treatment modality. We, hence, assessed the effects of LPCAT3 inhibitors, (R) HTS3 and HTS4, on sensitivity of the *KRAS^G12C^*-harboring MiaPaCa-2 cells to KRAS^G12C^ inhibitor Sotorasib. MiaPaCa-2 cells were treated with a fixed and sub-optimal 5 μM (R) HTS3 or HTS4 without/with varying concentrations of Sotorasib for 72 hours befor measuring cell viability. Combinations of Sotorasib and LPCAT3 inhibitors more effectively reduced viability of MiaPaCa-2 cells than Sotorasib alone (Fig.10A). We next treated MiaPaCa-2 cells with fixed and sub-optimal concentration of 2.5 nM of Sotorasib along with varying concentrations of (R) HTS3 and HTS4. Combinations of LPCAT3 inhibitors and Sotorasib more effectively reduced proliferation of MiaPaCa-2 cells than (R) HTS3 or HTS4 alone (Fig.10B). Taken together, LPCAT3 inhibitors enhance chemosensitivity of the KRAS-driven cancer cells to direct inhibition.

## Discussion

Spatial regulation of small GTPase has traditionally been understood at the level of organelle-scale compartmentalization. For instance, RAS isoforms and Rac1 predominantly function at the plasma membrane (PM), while Rabs function efficiently at various endomembranes. Recent super-resolution quantitative imaging, biophysical and computational studies identified a further tier of spatial organization, in which small GTPases further laterally segregate into nanometer-sized domains when localized to their favored compartments. This lateral spatial segregation is facilitated by their intricate ability to sense lipid headgroup and acyl chain structures, suggesting that functions of small GTPases depend on local membrane mesoscopic environments surrounding them and the metabolic alteration of lipid homeostasis in cells. Yet the molecular determinants and biological implications of the spatial sensitivity have been poorly understood. Here, we identify metabolic regulation of lipid acyl chain structures via LPCAT3 as a key mechanistic determinant of KRAS oncogenic signaling. These findings extend current models of KRAS compartmentalization beyond lipid headgroup specificity to include acyl chain composition as a key parameter and add another layer of regulation that transiently control small GTPase activities.

Our current understanding of lipid sensing by small GTPases predominantly centers on the interplay between membranes and their C- or N-terminal lipid anchors. The membrane-anchoring domains of many small GTPases contain long stretches of basic residues, implying extensive electrostatic interactions between small GTPases and lipids with anionic headgroups. Many small GTPases, such as KRAS, Rap1A and Rap1B, enrich PS lipids ^10,11,22,35^. Rac1 favors phosphoinositol 3,4,5-trisphosphate (PIP_3_) and phosphatidic acid (PA) ^36–38^. KRAS PM association is perturbed upon interference with distribution of PS lipids to the PM via vesicular trafficking of recycling endosomes ^39,40^, PM / endoplasmic reticulum (ER) contact sites ^41^ and interleaflet coupling with sphingolipids in the PM outer leaflet ^42^. Preference of KRAS for the monovalent PS lipid over multivalent phosphoinositol 4,5-bisphosphate (PIP_2_) and PIP_3_ suggests more than electrostatic contribution to lipid sensing of small GTPases. Indeed, shifting between the ordered and disordered conformations of KRAS lipid anchor, as well as equilibrium between allosteric orientation states of its G-domain, allow KRAS to differentiate nuanced structural differences of lipid headgroups and acyl chains and alter the KRAS preference for saturated and unsaturated PS lipids ^8,9,11^. Biologically, remodeling of lipid acyl chains, such as peroxidation of polyunsaturated acyl chains in ferroptosis ^31^ and modification of saturated/monounsaturated acyl chains via elongases ^43^, selectively modulates KRAS mutant activities. Our current findings further extent this model and highlight polyunsaturated lipids as critical components of the signaling nanoclusters of KRAS mutants, and targeting the polyunsaturated lipid-converting LPCAT3 selectively impacts oncogenic activities of KRAS mutants.

Ferroptosis is a new form of programmed cell death that is triggered by accumulation of peroxidized lipids. LPCAT3 is a critical upstream regulator of ferroptotic pathway because polyunsaturated lipids are more prone to peroxidation, thus serving as major substrates triggering ferroptosis. RAS signaling and activity are especially sensitive to induced ferroptosis. Various traditional ferroptosis inducers are named for their selective impact on RAS activity, such as Erastin (eradicator of RAS and Simian Virus 40 small T antigen-expressing cells ^44^) and RSL3 (Ras-Selective Lethal small molecule 3 ^45^). It is, thus, possible that effects of LPCAT3 on KRAS activity observed here may be linked to ferroptosis. Surprisingly, we observed that RSL3-induced ferroptosis has no effect on spatial distribution of KRAS^G12V^, despite enhancing PM localization and disrupting the clustering of PS lipids. This is different from how LPCAT3 downregulation disrupts PM association of KRAS^G12V^. These findings suggest that effects of LPCAT3 downregulation on KRAS mutant activity observed here are likely direct effects of polyunsaturated lipids on spatial signaling of KRAS mutants and unrelated to ferroptosis. This is supported by our findings that acute one-hour addback of unsaturated PS-enriched mouse brain PS extracts, but not saturated PS, restore the PM association of KRAS^G12V^. Thus, effects of LPCAT3 deletion on KRAS activity are likely results of changing polyunsaturated lipid homeostasis.

Since lipids are essential components in sustaining cell function, targeting lipid metabolism is usually perceived to cause high levels of cytotoxicity. Perturbations of lipid acyl chain homeostasis, and in turn the membrane core properties (such as elastivity and curvature), may also impact activities of all proteins sharing similar local membrane environments. However, recent studies are revealing selective sensitivity of KRAS mutant-driven tumor cells to perturbations of lipid acyl chain homeostasis, with minimal responses from cells expressing wild-type KRAS. For instance, KRAS mutant-driven tumor cells are especially more sensitive to accumulation of peroxidized lipids and the induced ferroptosis ^31,45^. Depletion of monounsaturated lipids via downregulation of elongase ELOVL6 also effectively abolishes oncogenic activities of KRAS-dependent tumor development ^43^. Perturbations of PS intracellular transport also demonstrate similar selective interference of KRAS oncogenesis ^39–42^. A possible explanation for this metabolic sensitivity is that these mutant KRAS-driven tumor cells reprogram their metabolic networks including lipid metabolism to sustain and fuel their KRAS-dependent pathological activities, thus becoming more addicted to the perturbed metabolism. Indeed, lipid profiles and enzymes within the lipid metabolic network differ significantly in cells expressing KRAS mutants and HRAS mutants ^42^, as well as those expressing KRAS mutants vs. wild-type KRAS ^43^. We recently also illustrated allele-specific lipid profiles, with KRAS^G12C^-expressing cells possessing higher levels of saturated lipids than KRAS^G12D^-expressing cells ^10^. This is consistent with our lipid mapping analysis showing that KRASG12D selectively enriches unsaturated lipids while KRASG12C gaining further enrichment of saturated lipids ^11^. Thus, targeting lipid metabolism may be more precise than previously thought. It is also possible that targeting the proteolipid signaling scaffolds shared by many KRAS mutants is a broad-spectrum strategy that simultaneously impacts multiple KRAS mutants.

## Conclusion

KRAS oncogenic signaling is predominantly compartmentalized to the proteolipid nanoclusters enriched with unsaturated lipids. We, here, propose that LPCAT3, by sustaining homeostasis of unsaturated lipids, functions as a regulator of membrane biophysical state that determines the capacity of KRAS to organize into signaling-competent nanostructures. By coupling lipid metabolism to spatial and allosteric regulation of KRAS, LPCAT3 defines a previously underappreciated vulnerability in oncogenic signaling—one that can be exploited in a mutation-agnostic manner across KRAS-driven cancers.

## Acknowledgements

This work was supported in part by the National Institutes of Health R01GM138668 to N. Arora, H. Liang and Y. Zhou. Y Zhou is also in part supported by the U Foundation Pilot Award.

## Materials and Methods

### Cell line maintenance

MiaPaCa-2, BxPC3, and MOH were kindly provided by Dr Craig Logsdon at MD Anderson Cancer, Houston, TX. AsPC1, Patu8902S, Capan-1, iKRAS AK192 were kindly provided by Dr Haoqing Ying at MD Anderson Cancer, Houston, TX. TLA293T cells were a generous gift from Dr Guangwei Du, McGovern Medical School, Houston, TX. MiaPaCa-2 cells were maintained in DMEM media supplemented with 2 mM L-glutamine, 10% FBS and 2.5% horse serum. MiaPaCa-2 vector control and LPCAT3 knockdown cells (LPCAT3KD) were maintained in DMEM media supplemented with 2 mM L-glutamine, 10% FBS and 2.5% horse serum, 1µg/mL puromycin. BxPC-3 and MOH cells were grown in RPMI media supplemented with 10% FBS. HPNE cells were cultured in 75% DMEM and 25% Medium M3 Base supplemented with 5% FBS, 10 ng/mL human recombinant EGF, 5.5 mM D-glucose, and 750 ng/mL puromycin. iKRAS AK192 cells were maintained in RMPI media supplemented with Tetracycline Negative 10% FBS and 1ug/mL doxyclycine. All other cell lines were maintained in DMEM supplemented with 10% FBS. All the cells were grown in 5% CO_2_ incubator at 37°C.

### Generation of CRISPR/Cas9 cell lines

MiaPaca-2 and BxPC-3 cells were transduced with *OSBPL8* sgRNA (3′-TGCAAATCTTTGGTTGGCGT-5′) plus Cas9 expression followed by puromycin selection (3 μg/mL) after 48 h. Single colonies were generated from the pool of polyclonal knockdown cells.

### Total RNA extraction and sequencing

Total RNA extraction followed the QiagenMag-B ind total RNA Xpress kit protocol. MiaPaCa-2 cells containing vector control or *LPCAT3KD were* grown to confluency were trypsinized and lysates were combined with Mag-Bind^®^ Particles RQ, vortexed and magnetically separated in the presence of isopropanol. The bound genomic DNA was digested with DNase I directly on the magnetic particles. Beads were then washed with alcohol, and the pure RNA was eluted in nuclease-free water for analysis.

Total RNA quality was assessed by Novogene prior to library preparation. Poly(A) mRNA libraries were prepared and sequenced by Novogene (Sacramento, California, USA) as 150 bp paired-end reads on an Illumina NovaSeq X Plus. Raw data (raw reads) was first processed through fastp software to clean reads, remove adapters, reads containing poly-N and low-quality reads. All the downstream analyses were based on the clean data with high quality. Then, processed reads were aligned to the reference genome (hg38) using Hisat2 v2.0.5 and featureCounts v1.5.0 was used to count the reads numbers mapped to each gene. Then Fragments Per Kilobase of transcript sequence per Millions base pairs sequenced (FPKM) of each gene were calculated based on the length of the gene and reads count mapped to this gene. For differential expression analyses DESeq2 for two conditions (control vs. knocked down) was performed and genes with fold change (FC) > 1.5 (or Log2Fold Change > 0.58) and FDR <0.05 were assigned as differentially expressed genes (DEG). P-values were adjusted using the Benjamini-Hochberg (BH) procedure. Then, standard gene ontology analyses were performed with package clusterProfiler.

### Lipidomics

MiaPaCa-2 vector control and *LPCAT3*KD cells were grown to confluency, washed with PBS and trypsinized using 1X trypsin-EDTA. Following centrifugation, cells were washed twice with ice-cold PBS, resuspended in PBS for subsequent cell number counting via a Countess automated cell counter (Invitrogen). Final concentration was 450,000 cells / 300 μL PBS for all samples, which were frozen at −80°C and handed to the Metabolomics core at the MD Anderson Cancer Center (Houston, Texas, USA) for detailed lipidomics analysis. Three independent experiments were performed and analyzed. Data are shown as mean ± SEM. Statistical significance was evaluated using one-way ANOVA, with ^∗^ indicating p < 0.05.

### EM Spatial Analysis

#### Univariate nanoclustering analysis quantifies spatial distribution of a single species on the native cell plasma membrane

Human pancreatic ductal adenocarcinoma (PDAC) cell line MiaPaCa-2 vector control and *LPCAT3*KD cells were maintained in Dulbecco’s modified Eagle’s medium (DMEM) containing 10% fetal bovine serum (FBS) and 2.5% horse serum. Cells were seeded on gold EM grids to grow overnight, followed by preparation of the cell plasma membrane (PM) for electron microscopy (EM) analysis.

EM spatial distribution method is used to quantify the extent of KRAS protein lateral spatial segregation in the inner leaflet of the PM. Briefly, MiaPaCa-2 cells expressing GFP-tagged KRAS^G12V^mutant were grown on pioloform and poly-L-lysine coated gold grids. The gold grids were washed in PBS to remove medium and then a PBS-soaked Whatman filter paper was applied for 5 min for removal of apical PM. Basal PM on the gold grids were then fixed in fixing reagent (4% paraformaldehyde and 0.1% glutaraldehyde dissolved in potassium acetate (KOAc) buffer), labelled with 4.5 nm gold particles coupled to anti-GFP antibody, and stained with 0.3% uranyl acetate. The grids were scooped in EM loops and dried overnight prior to imaging.

Distribution of gold particles on the basal PM sheets was imaged using JEOL JEM-1400 transmission EM at 100,000x magnification. The EM images were analyzed using ImageJ software to assign x and y coordinates to gold particles in a 1 μm^2^ area of interest on the PM sheets. We use Ripley’s K-function to quantify the distribution and the extent of nanoclustering of gold-labeled GFP-RAS on the PM (Equation A and B).

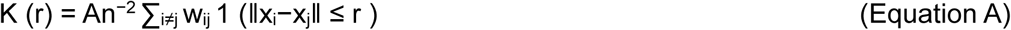

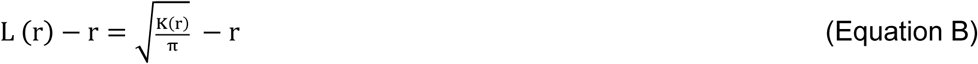

where K(r) is the univariate K-function for n gold particles in the area A; r is radius; || **·** || is Euclidean distance; 1(**·**) is the indicator function that has a value of 1 if ||x_i_-x_j_|| ≤ r and 0 otherwise; and w_ij_^-1^ is the proportion of the circumference of a circle with center at x_i_ and a radius ||x_i_-x_j_|| located within A. *L*(*r*) - *r* is a linear transformation of *K*(*r*) and is normalized on the 99% confidence interval (99% C.I.) estimated from Monte Carlo simulations. The maximum value of the *L*(*r*) - *r* function: L_max_ correlates with the extent of clustering. For each treatment condition, at least 15 PM sheets were imaged, analyzed and the data pooled. Statistical significance was evaluated using Bootstrap tests as previously described^46,47^.

#### Bivariate co-clustering analysis quantifies co-localization between two different species on the native cell PM

Similar to the univariate analysis described above, in bivariate analyses the PM sheets of MiaPaCa-2 cells co-expressing GFP-LactC2 (probing PS lipids) and RFP-tagged KRAS^G12V^ were attached to EM grids and fixed with 4% paraformaldehyde and 0.1% glutaraldehyde and stained sequentially with anti-GFP 6-nm gold and anti-RFP 2-nm gold conjugated antibodies before imaged and analyzed. A bivariate K-function analysis tested the null hypothesis that the two populations of gold particles spatially separate from each other, were calculated with bivariate K-functions (Equations C, D, E and F).

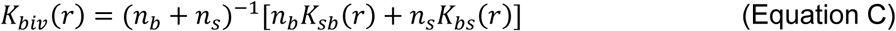

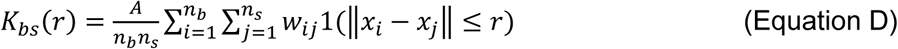

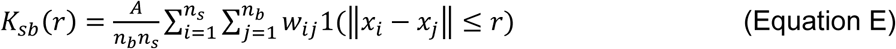

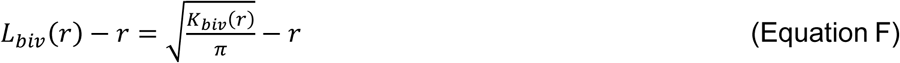

Where, *K_biv_*(*r*) denotes a bivariate estimator and contains two individual bivariate K-functions: *K_bs_*(*r*) quantifies how the big 6 nm gold particles (*b* = big gold) distribute around each 2 nm small gold particle (*s* = small gold); *K_sb_*(*r*) describes how small gold particles distribute around each big gold particle. The value of n_b_ indicates the number of 6 nm big gold and n_s_ indicates the number of 2nm small gold within a PM area of *A*. Other parameters denote the same definitions as defined in the univariate calculations in Eqs.A and B. *L_biv_*(*r*)-*r* is a linear transformation of *K_biv_*(*r*) and is normalized against the 95% confidence interval (95% C.I.). An *L_biv_*(*r*)-*r* value of 0 indicates spatial segregation between the two populations of gold particles, whereas an *L_biv_*(*r*)-*r* value above the 95% C.I. of 1 at the corresponding distance of *r* indicates yields statistically significant co-localization at certain distance yields. Area-under-the-curve for each *L_biv_*(*r*)-*r* curves were calculated within a fixed range 10 < *r* < 110 nm, and was termed bivariate *L_biv_*(*r*)-*r* integrated (or LBI):

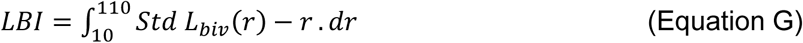

For each treatment condition, at least 15 PM sheets were imaged, analyzed and pooled, shown as mean of LBI values ± SEM. Statistical significance was evaluated using Bootstrap tests as previously described^46,47^.

### Western blotting

Human and murine pancreatic cancer cell lines were grown to confluency. For siRNA and LPCAT3 inhibitors treatment, cells were treated for 48 h followed by protein isolation for expression analysis. Cells were washed twice with ice cold PBS followed by incubation on ice for 10 min with lysis buffer containing 50 mM Tris (pH 7.5), 1% NP40, 75 mM NaCl, 5 mM MgCl_2_, 5 mM EGTA, 25 mM NaF, 0.5 μg/mL aproptinin, 0.1 mM Na_3_VO_4_, 3 μg/mL Leupeptin, 1 mM DTT. Cell lysate was collected and centrifuged at 1000 RPM at 4°C for 5 min. Supernatant was collected and concentration of total protein was measured by Bradford assay. 10-20 μg of total protein was mixed with sample buffer containing 1 mM Tris (pH6.8), 20 mg/mL SDS, 10% glycerol, 0.5 mg/mL bromophenol blue dye and 15.4 mg/mL DTT and heated at 95°C for 5 min before being resolved by electrophoresis in a 10% SDS-polyacrylamide gel. Proteins were electro-transferred to polyvinylidene difluoride (PVDF) membrane using pre-chilled transfer buffer. PVDF membranes were rinsed twice with TBST (10 mM Tris (pH 7.5), 150 mM NaCl and 0.1% Tween 20), for 5 min and membrane were blocked for 1 hr in 3% BSA dissolved in TBST for phospho-antibodies or 5% skim milk in TBST for other proteins. Membranes were incubated overnight in primary antibody in blocking solution. The PVDF membrane was then washed thrice with TBST (10 min each) before a 1-h incubation with secondary antibody in TBST buffer containing 5% skim milk: anti-rabbit/anti-mouse at 1/5000 dilution. PVDF membranes were washed thrice in TBST (10 min each) before a 2 min incubation with enhanced chemiluminescence (ECL) solution. At least 3 independent experiments were conducted. Data are shown as mean ± SEM. ImageJ software analysis was used to evaluate expression intensity and identify fold change. Statistical significance was evaluated using one-way ANOVA with ^∗^ indicating p < 0.05.

### siRNA mediated knockdown of LPCAT3

siRNAs targeting murine LPCAT3 were procured from Sigma Aldrich (Product Number SASI_Mm02_00340517, SASI_Mm01_00197994, SASI_Mm01_00197995). iKRAS AK192 cells were maintained in RMPI media supplemented with Tetracycline Negative 10% FBS and 1ug/mL doxyclycine. Cells were seeded at a density of 7×10^4^ in 12-well tissue culture plates and incubated in CO_2_ incubator overnight. Following confluency, cells were transfected with siRNA at a concentration of 10 nM and grown in OPTI-MEM media for 48 h prior to harvesting for cellular analysis.

### Proliferation and Colony formation Assay

For proliferation assay, MiaPaCa-2 vector control and *LPCAT3*KD (1×10^3^) cells were seeded in 96-well tissue culture plates. Cells were allowed to grow for another 72h followed by quantification of cell number. LPCAT3 inhibitor studies, MiaPaCA-2 (1×10^3^), MOH (1×10^3^), AsPC1 (1×10^3^), Patu(1×10^3^), Capan-1(1×10^3^) and BxPC-3(3×10^3^) cells were seeded in 96-well plate. The following day fresh growth media containing 1% DMSO or different inhibitor concentrations was added to the cells and treated for 72h. For combination therapy MiaPaCa-2 (1×10^3^) cells were seeded in 96-well tissue culture plates. After 24h, cells were treated with varying concentrations of LPCAT3 inhibitors (0.5-5 µM) and fixed concentration of KRAS^G12C^ inhibitor Sotorasib (2.5 nM) and treated for 48 h. Cell numbers were determined by CyQuant Proliferation Assay (ThermoFisher Scientific) according to the manufacturer’s protocol.

For colony formation assay, MiaPaCa-2 vector control and *LPCAT3*KD (100 cells/well) and MiaPaCa-2 (100 cells/well) cells were seeded in 6-well tissue culture plate for 10 days and iKRAS AK192 cells (250 cells/well) were seeded in 6-well tissue culture plate for a week. For LPCAT3 inhibitor study, cells were treated with 10 µM of HTS-3 or HTS-4. Following treatment, cells were stained to evaluate colony number. Briefly, cells were washed twice with PBS, followed by fixing with 4% paraformaldehyde for 15 min. cell stating was performed with 0.01% crystal violet for 15 min. Colony images were captured using Perkin Elmer X3 multiplate reader. Colony count was performed using ImageJ.

### Chemicals and reagents

| REAGENTS | SOURCE | CATALOG NUMBER |
| --- | --- | --- |
| DMEM | CYTIVA | SH30243.01 |
| FBS | GIBCO | 16000-044 |
| Dialyzed FBS | GIBCO | 26400-036 |
| BCS | Hyclone | SH30072.03 |
| Trypsin-EDTA (0.5%) | GIBCO | 15400-054 |
| Trypsin-EDTA (0.25%) | GIBCO | 25200-072 |
| Lipofectamine | Invitrogen | 50470 |
| Opti-MEM | GIBCO | 31985-070 |
| Pioloform | TED PELLA | 19244 |
| Choroform | Electron Microscopy<br>Sciences (EMS) | 12550 |
| Poly-L-lysine | Sigma | P8920-100ML |
| EM copper grids | EMS | G200-CU |
| Trisodium citrate | Sigma | C3674-100G |
| Tannic acid | EMS | 21710 |
| Potassium carbonate | Sigma | P5833-500G |
| Potassium acetate | Sigma | P1190-1KG |
| Glycerol | Sigma | G55-1L |
| Paraformaldehyde | EMS | 1570 |
| Glutaraldehyde | EMS | 16220 |
| Fish gelatin | Sigma | G7765-250ML |
| BSA | Sigma | A6003-100G |
| Gold chloride | Sigma | 254169-5G |

