## Supplementary figures and images for "Lipid acyl chain remodeling governs KRAS nanoclustering and oncogenic functions through membrane composition control"

### supplementary figures

**Supplemental Figure 1**

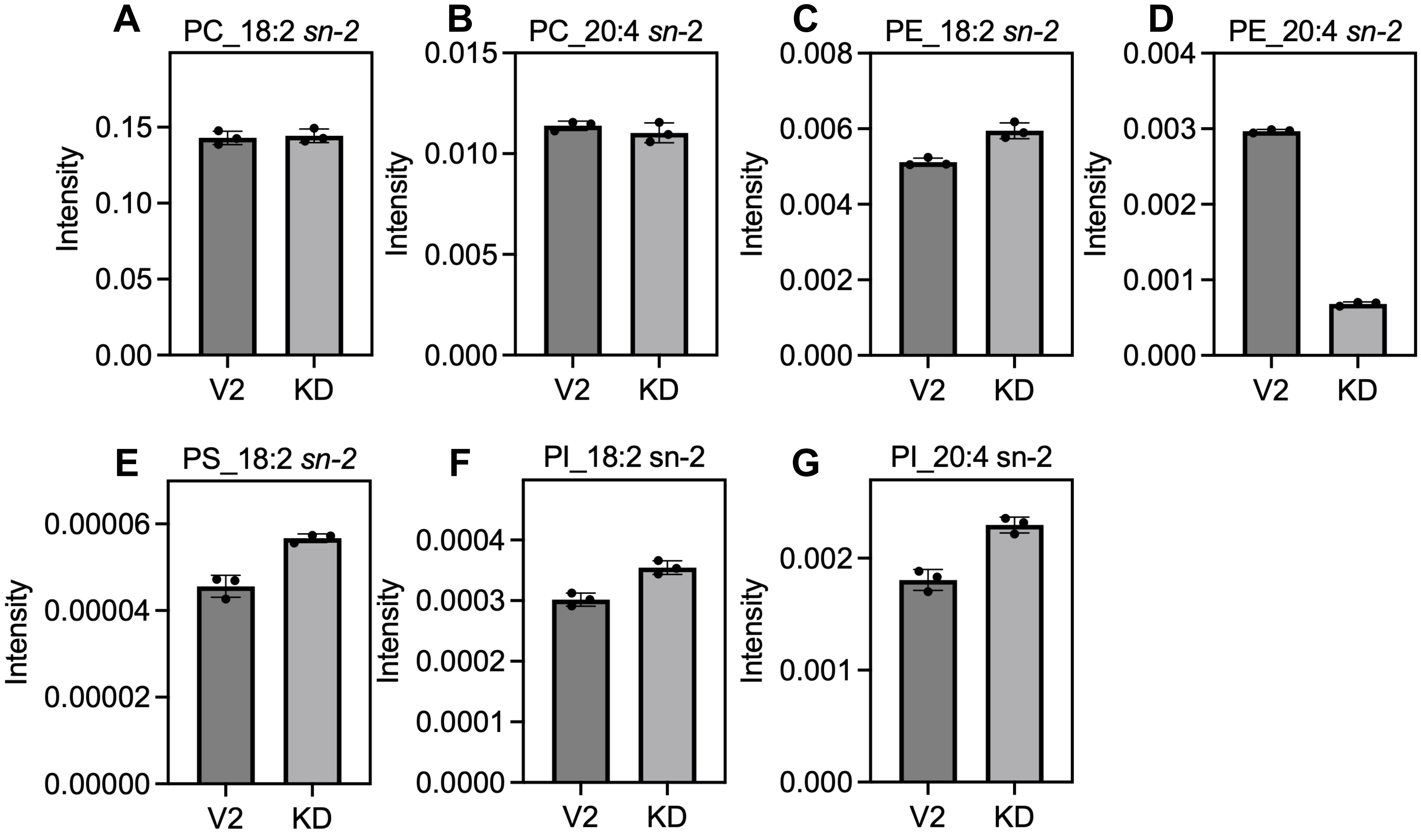

#### Supplemental Figure 2

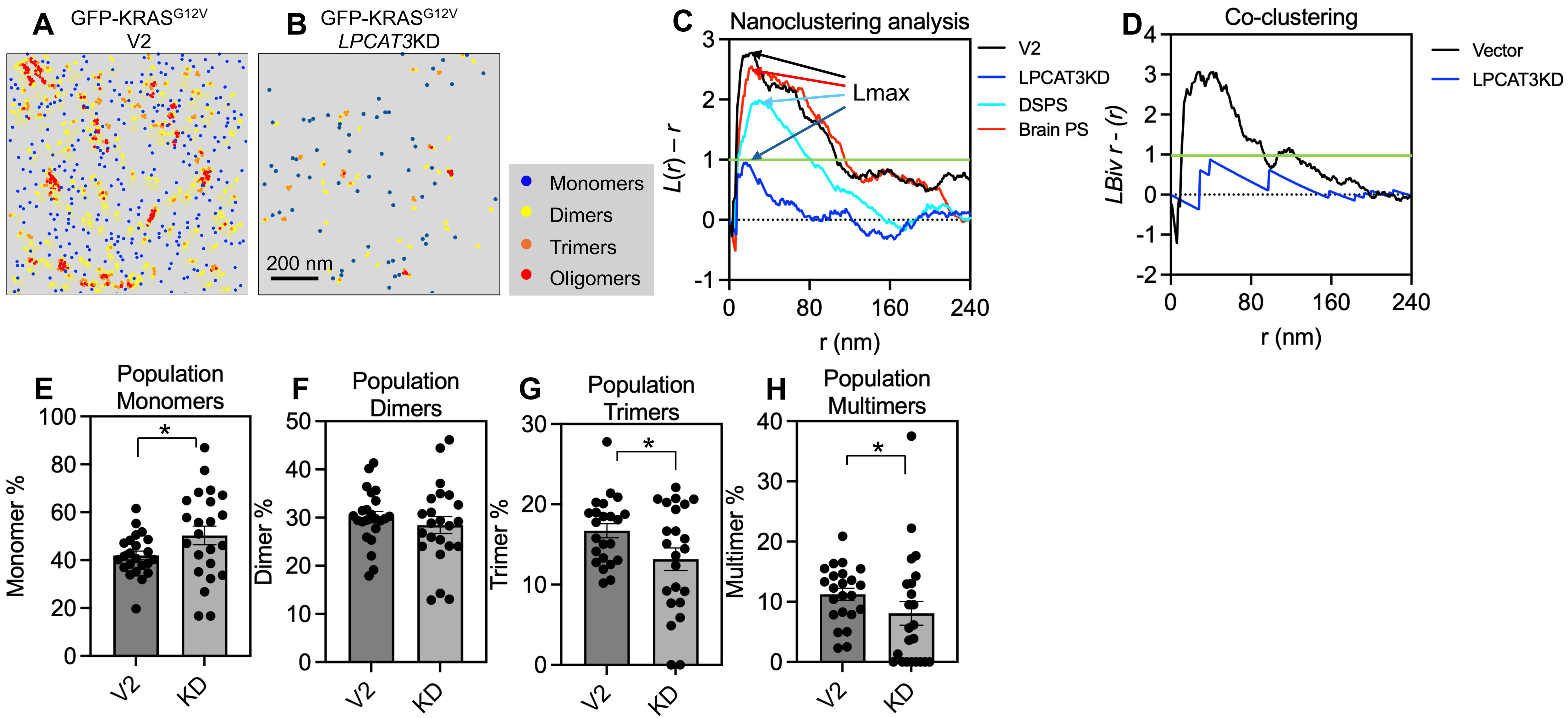

**Supplemental Figure 3**

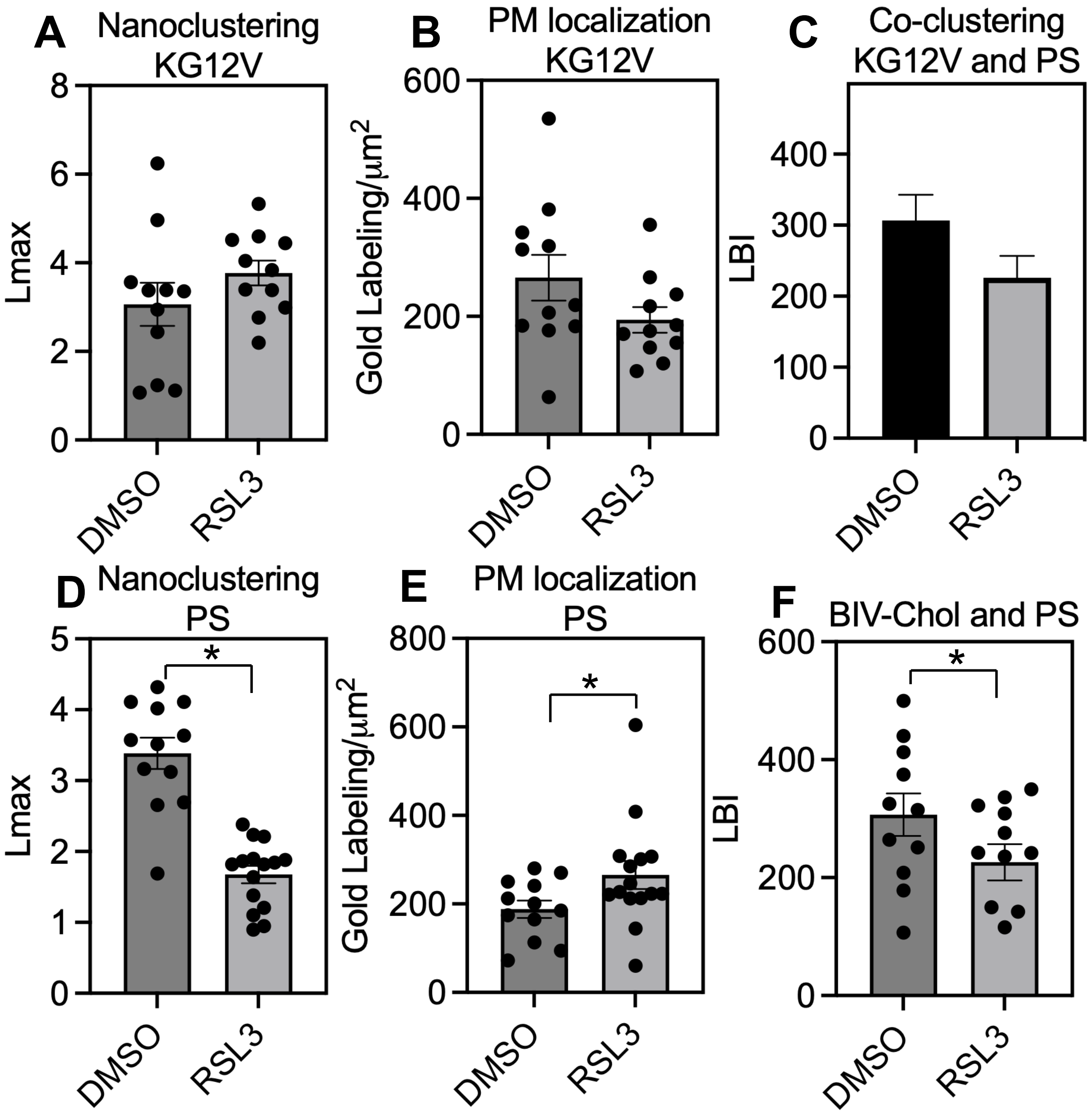
